# Convergent recruitment of life cycle regulators to direct sporophyte development in two eukaryotic supergroups

**DOI:** 10.1101/460436

**Authors:** Alok Arun, Susana M. Coelho, Akira F. Peters, Simon Bourdareau, Laurent Peres, Delphine Scornet, Martina Strittmatter, Agnieszka P. Lipinska, Haiqin Yao, Olivier Godfroy, Gabriel J. Montecinos, Komlan Avia, Nicolas Macaisne, Christelle Troadec, Abdelhafid Bendahmane, J. Mark Cock

## Abstract

Three amino acid loop extension homeodomain transcription factors (TALE HD TFs) act as life cycle regulators in green algae and land plants. In mosses these regulators are required for the deployment of the sporophyte developmental program. We demonstrate that mutations in either of two TALE HD TF genes, *OUROBOROS* or *SAMSARA*, in the brown alga *Ectocarpus* result in conversion of the sporophyte generation into a gametophyte. The OUROBOROS and SAMSARA proteins heterodimerise in a similar manner to TALE HD TF life cycle regulators in the green lineage. These observations demonstrate that TALE-HD-TF-based life cycle regulation systems have an extremely ancient origin, and that these systems have been independently recruited to regulate sporophyte developmental programs in at least two different complex multicellular eukaryotic supergroups, Archaeplastida and Chromalveolata.

## Introduction

Developmental processes need to be precisely coordinated with life cycle progression. This is particularly important in multicellular organisms with haploid-diploid life cycles, where two different developmental programs, corresponding to the sporophyte and gametophyte, need to be deployed appropriately at different time points within a single life cycle. In the unicellular green alga *Chlamydomonas,* plus and minus gametes express two different HD TFs of the three amino acid loop extension (TALE) family called Gsm1 and Gsp1 (Lee et al., 2008). When two gametes fuse to form a zygote, these two proteins heterodimerise and move to the nucleus, where they orchestrate the diploid phase of the life cycle. Gsm1 and Gsp1 belong to the knotted-like homeobox (KNOX) and BEL TALE HD TF classes, respectively. In the multicellular moss *Physcomitrella patens*, deletion of two KNOX genes, *MKN1* and *MKN6*, blocks initiation of the sporophyte program leading to conversion of this generation of the life cycle into a diploid gametophyte (Sakakibara et al., 2013). Similarly, the moss BEL class gene *BELL1* is required for induction of the sporophyte developmental program and ectopic expression of *BELL1* in gametophytic tissues induces the development of apogametic sporophytes during the gametophyte generation of the life cycle (Horst et al., 2016). In mosses, therefore, the KNOX and BEL class life cycle regulators have been recruited to act as master regulators of the sporophyte developmental program, coupling the deployment of this program with life cycle progression. *P. patens* KNOX and BEL proteins have been shown to form heterodimers (Horst et al., 2016) and it is therefore possible that life cycle regulation also involves KNOX/BEL heterodimers in this species.

The filamentous alga *Ectocarpus* has emerged as a model system for the brown algae (Cock et al., 2015; Coelho et al., 2012). This alga has a haploid-diploid life cycle that involves alternation between multicellular sporophyte and gametophyte generations (Figure 1A). A mutation at the *OUROBOROS* (*ORO*) locus has been shown to cause the sporophyte generation to be converted into a fully functional (gamete-producing) gametophyte (Figure 1B) (Coelho et al., 2011). This mutation therefore induces a phenotype that is essentially identical to that observed with the *P. patens mkn1 mkn6* double mutant, but in an organism from a distinct eukaryotic supergroup (the stramenopiles), which diverged from the green lineage over a billion years ago (Eme et al., 2014).

**Figure 1.**
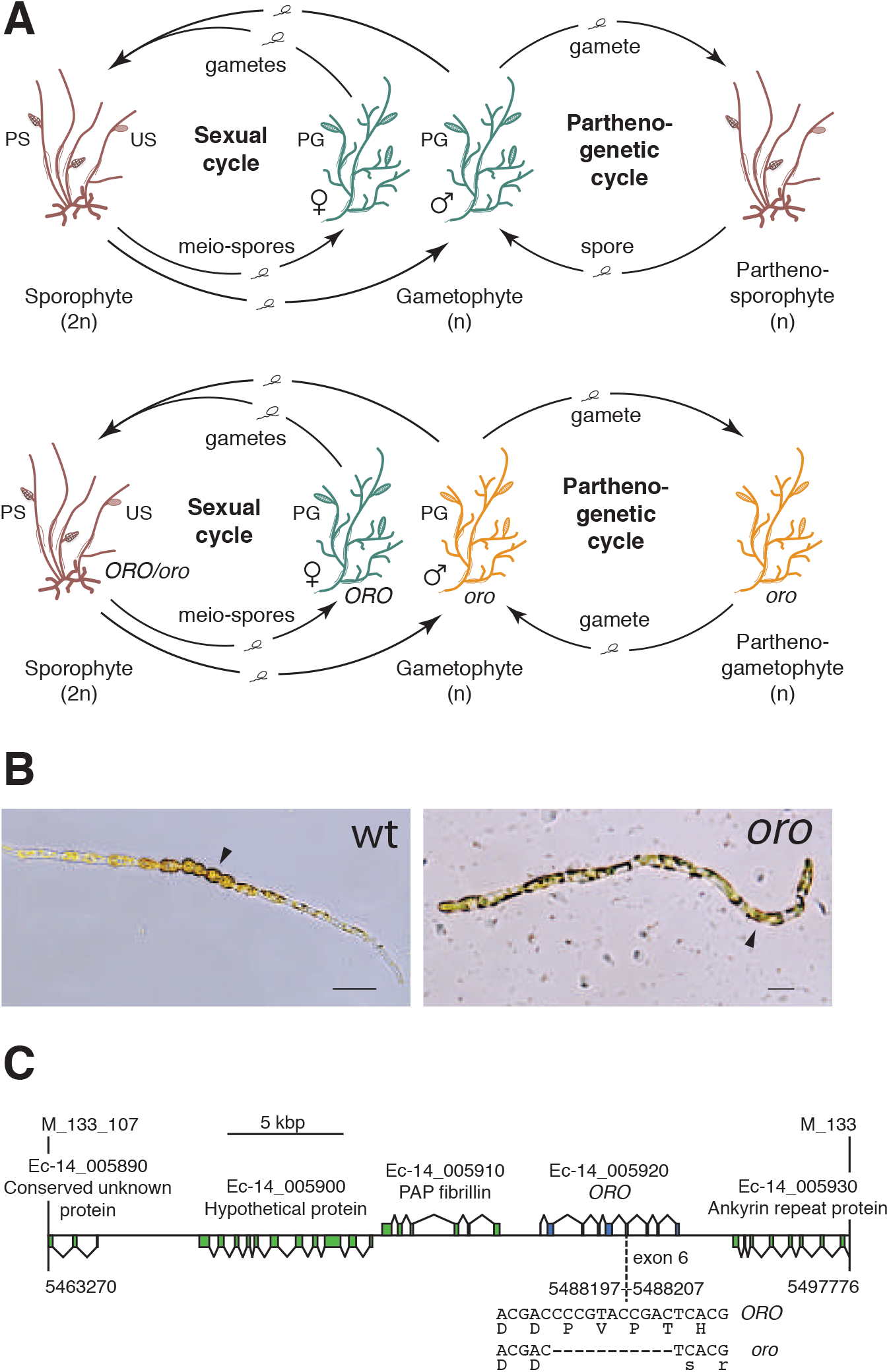
The *oro* life cycle mutation corresponds to a TALE homeodomain transcription factor gene. **(A)** Life cycle of wild type and *oro* mutant *Ectocarpus*. The wild type sexual cycle (upper panel) involves production of meio-spores by the diploid sporophyte via meiosis in unilocular (single-chambered) sporangia (US). The meio-spores develop as haploid, dioecious (male and female) gametophytes. The gametophytes produce gametes in plurilocular gametangia (PG), which fuse to produce a diploid sporophyte. Gametes that fail to fuse can develop parthenogenetically to produce a partheno-sporophyte, which can produce spores by apomeiosis or following endoreduplication to engender a new generation of gametophytes. PS, plurilocular sporangium (asexual reproduction). Gametes of the *oro* mutant (lower panel) are unable to initiate the sporophyte program and develop parthenogenetically to produce partheno-gametophytes. The mutation is recessive so a cross with a wild type gametophyte produces diploid sporophytes with a wild type phenotype. **(B)** Young gamete-derived parthenotes of wild type and *oro* strains. Arrowheads indicate round, thick-walled cells typical of the sporophyte for the wild type and long, wavy cells typical of the gametophyte for the *oro* mutant. Scale bars: 20 μm. **(C)** Representation of the interval on chromosome 14 between the closest recombining markers to the *ORO* locus (M_133_107 and M_133) showing the position of the single mutation within the mapped interval.

Here we identify mutations at a second locus, *SAMSARA*, that also result in conversion of the sporophyte generation into a gametophyte. Remarkably, both *OUROBOROS* and *SAMSARA* encode TALE HD TFs and the two proteins associate to form a heterodimer. These observations indicate that TALE-HD-TF-based life cycle regulatory systems have very deep evolutionary origins and that they have been independently recruited in at least two eukaryotic supergroups to act as master regulators of sporophyte developmental programs.

## Results

### Two TALE homeodomain transcription factors direct sporophyte development

The *ORO* gene was mapped to a 34.5 kbp (0.45 cM) interval on chromosome 14 using a segregating family of 2000 siblings derived from an *ORO* x *oro* cross and a combination of amplified fragment length polymorphism (AFLP) (Vos et al., 1995) and microsatellite markers. Resequencing of the 34.5 kbp interval in the *oro* mutant showed that it contained only one mutation: an 11 bp deletion in exon six of the gene with the LocusID Ec-14_005920, which encodes a TALE homeodomain transcription factor. (Figure 1C).

A visual screen of about 14,000 UV-mutagenised germlings identified three additional life cycle mutants (designated *samsara-1, samsara-2* and *samsara-3,* abbreviated as *sam-1, sam-2* and *sam-3*). The *sam* mutants closely resembled the *oro* mutant in that gamete-derived parthenotes did not adopt the normal sporophyte pattern of development but rather resembled gametophytes. Young, germinating individuals exhibited the wavy pattern of filament growth typical of the gametophyte and, at maturity, never produced unilocular sporangia (the reproductive structures where meiosis occurs; Figure 1A), a structure that is uniquely observed during the sporophyte generation (Figure 2A-C-figure supplement 1). Moreover, the *sam* mutants exhibited a stronger negative phototrophic response to unilateral light than wild type sporophytes (Figure 2D), a feature typical of gametophytes (Peters et al., 2008) that was also observed for the *oro* mutant (Coelho et al., 2011).

**Figure 2.**
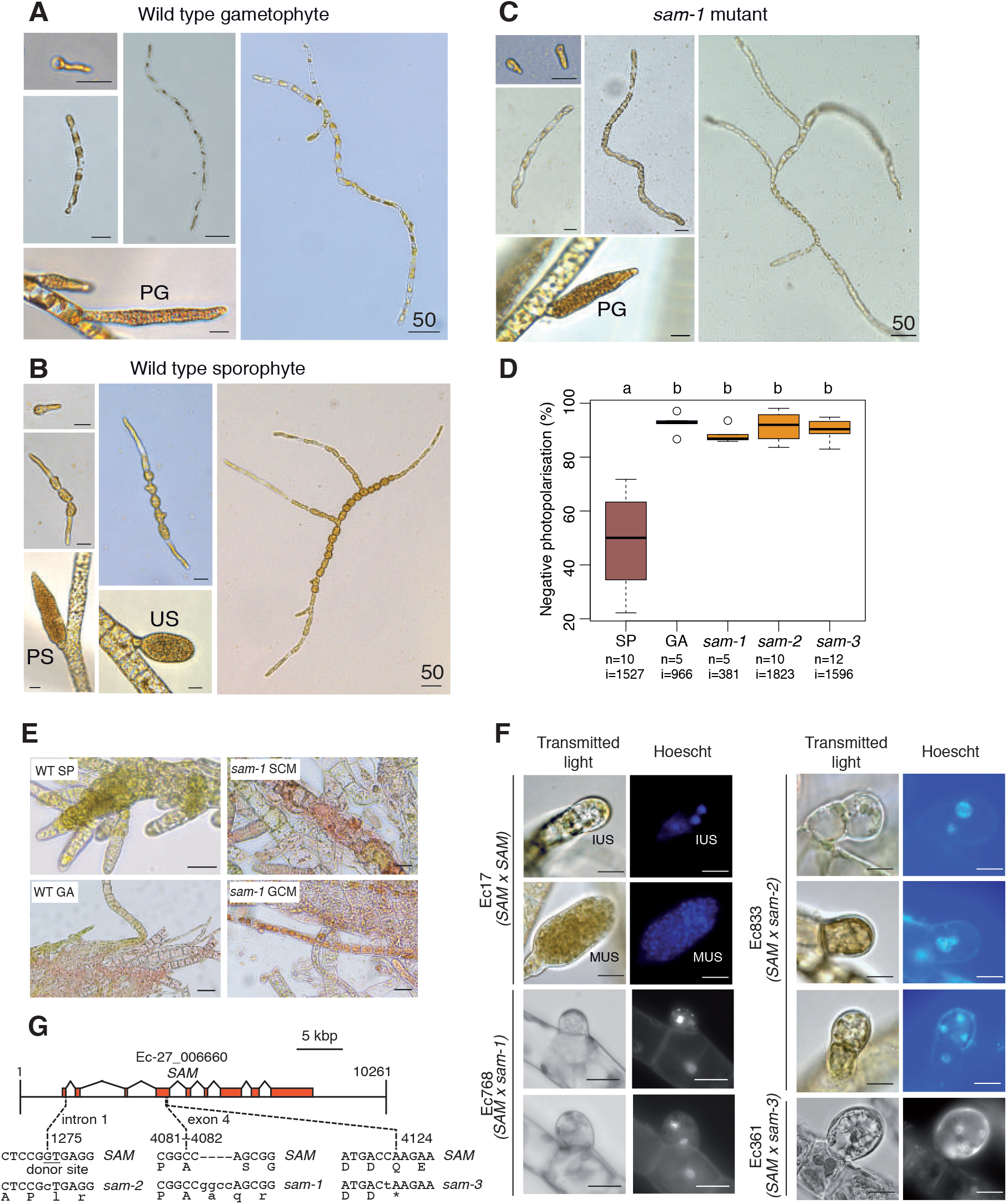
Phenotypic and genetic characterisation of *sam* life cycle mutants. **(A-C)** The *sam-1* mutant exhibits gametophyte-like morphological characteristics. Different stages of (A) wild type gametophyte (strain Ec32), (B) wild type partheno-sporophyte (strain Ec32) and (C) *sam-1* mutant (strain Ec374). PG, plurilocular gametangia; PS, plurilocular sporangium; US, unilocular sporangium. **(D)** *sam* mutants exhibit a gametophyte-like photopolarisation response to unidirectional light. Letters above the boxplot indicate significant differences (Wilcoxon test, p-value<0.01). n, number of replicates; i, number of individuals scored. **(E)** Representative images of congo red staining showing that the *sam-1* mutant protoplasts are resistant to treatment with sporophyte conditioned medium (SCM). GCM, control gametophyte conditioned medium. **(F)** Abortion of unilocular sporangia in *sam-1, sam-2* or *sam-3* mutant sporophytes. Images are representative of n=19 (Ec17), n=23 (Ec768), n=20 (Ec833) and n=14 (Ec361) unilocular sporangia. IUS, immature unilocular sporangium; MUS, mature unilocular sporangium. **(G)** Locations of the three *sam* mutations within the *SAM* gene. Scale bars: 20 μm (or 50μm if indicated by 50).

Genetic crosses confirmed that the *sam* mutants were fully functional (i.e. gamete-producing) gametophytes and complementation analysis indicated that they were not located at the same genetic locus as the *oro* mutation (Table supplement 1). Interestingly, hybrid sporophytes that were heterozygous for the *sam* mutations failed to produce functional unilocular sporangia. Wild type unilocular sporangia contain about a hundred haploid meio-spores produced by a single meiotic division followed by several rounds of mitotic divisions, whereas unilocular sporangia of *SAM/sam* heterozygotes never contained more than four nuclei indicating that abortion was either concomitant with or closely followed meiosis (Figure 2F). This indicated either a dominant effect of the *sam* mutations in the fertile sporophyte or abortion of the sporangia due to arrested development of the two (haploid) meiotic daughter cells that carried the mutant *sam* allele.

*Ectocarpus* sporophytes produce a diffusible factor that induces gametophyte initial cells or protoplasts of mature gametophyte cells to switch to the sporophyte developmental program (Arun et al., 2013). The *oro* mutant is not susceptible to this diffusible factor (*oro* protoplasts regenerate as gametophytes in sporophyte-conditioned medium) indicating that *ORO* is required for the diffusible factor to direct deployment of the sporophyte developmental pathway (Arun et al., 2013). We show here that the *sam-1* mutant is also resistant to the action of the diffusible factor. Congo red staining of individuals regenerated from *sam-1* protoplasts that had been treated with the diffusible factor detected no sporophytes, whereas control treatment of wild type gametophyte-derived protoplasts resulted in the conversion of 7.5% of individuals into sporophytes (Figure 2E-table supplement 2). Therefore, in order to respond to the diffusible factor, cells must possess functional alleles of both *ORO* and *SAM*.

The *Ectocarpus* genome contains two TALE HD TFs in addition to the *ORO* gene. Resequencing of these genes in the three *sam* mutants identified three genetic mutations, all of which were predicted to severely affect the function of Ec-27_006660 (Figure 2G). The identification of three disruptive mutations in the same gene in the three independent *sam* mutants strongly indicates that these are the causative lesions. Ec-27_006660 was therefore given the gene name *SAMSARA (SAM)*. *ORO* and *SAM* transcripts were most abundant in gametes (Figure 3A), consistent with a role in initiating sporophyte development following gamete fusion. Quantitative PCR experiments demonstrated that sporophyte and gametophyte marker genes (Peters et al., 2008) were down- and up-regulated, respectively, in *sam* mutant lines (Figure 3B), as was previously demonstrated for the *oro* mutant (Coelho et al., 2011).

**Figure 3.**
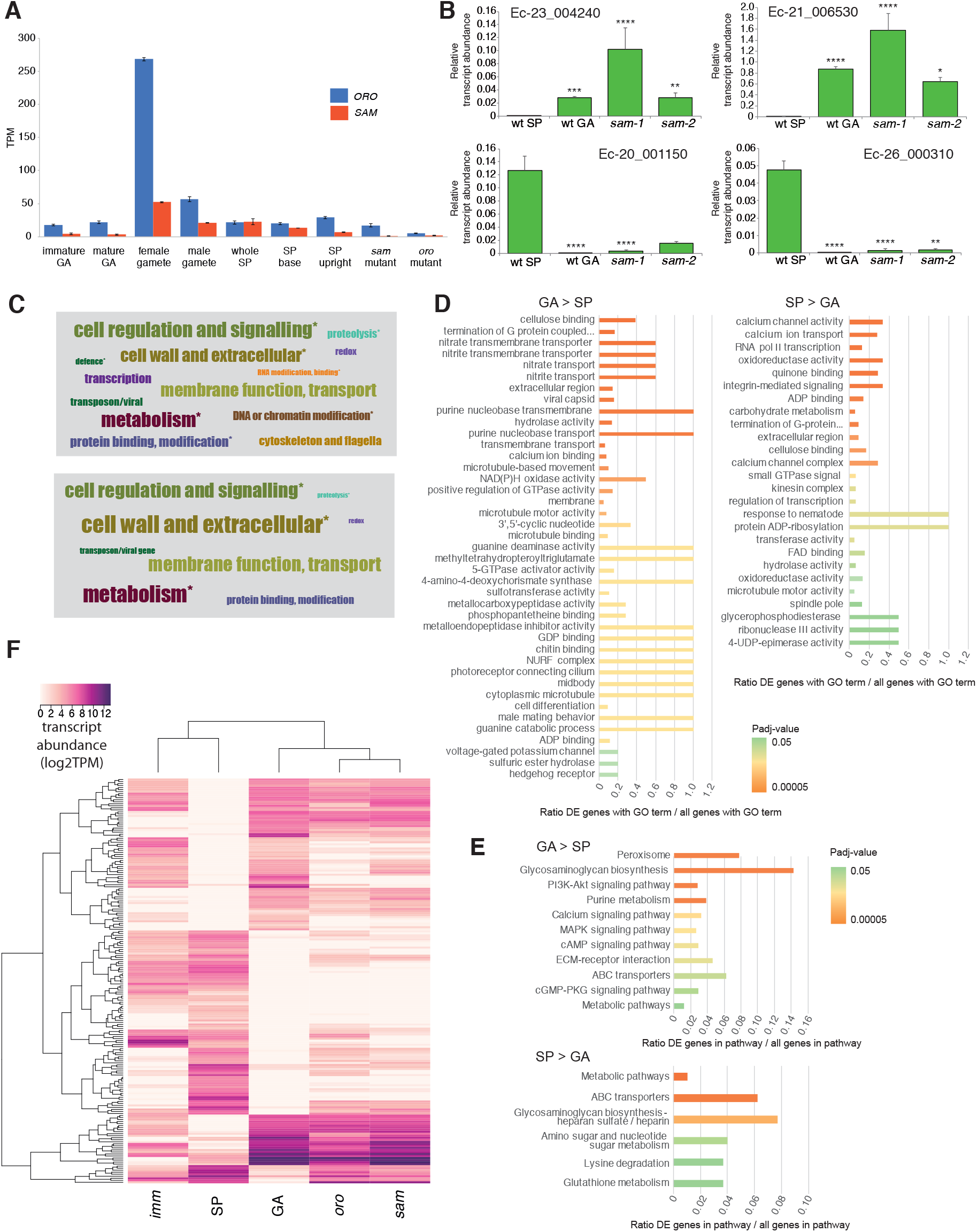
Gene expression analysis. **(A)** Abundance of *ORO* and *SAM* transcripts during different stages of the life cycle. Error bars, standard error of the mean (SEM); TPM: transcripts per million. **(B)** Quantitative reverse transcription PCR analysis of generation marker genes. The graphs indicate mean values ± standard error of transcript abundances for two gametophyte marker genes, Ec-23_004240 and Ec-21_006530, and two sporophyte marker genes, Ec-20_001150 and Ec-26_000310. Data from five independent experiments. * p ≤ 0.05, ** p ≤ 0.01, *** p ≤ 0.001, **** p ≤ 0.0001. **(C)** Word cloud representations of the relative abundances (log2 gene number) of manually assigned functional categories in the set of genes that were differential regulated between the sporophyte and gametophyte generations (upper panel) and in the subset of those genes that encode secreted proteins (lower panel). Asterisks indicate functional categories that were significantly over- or under-represented in the two datasets. **(D-E)** Significantly overrepresented GO terms (D) and KEGG pathways (E) associated with generation-biased genes. **(F)** Expression patterns of the 200 genes most strongly generation-biased genes. *oro*, *oro* mutant; *sam*, *sam* mutant; *imm*, *immediate upright* mutant; GA: gametophyte; SP: sporophyte.

### *ORO* and *SAM* regulate the expression of sporophyte generation genes

To investigate the genetic mechanisms underlying the switch from the gametophyte to the sporophyte program directed by the *ORO* and *SAM* genes, we characterised the gene expression networks associated with the two generations of the *Ectocarpus* life cycle. Comparative analysis of sporophyte and gametophyte RNA-seq data identified 1167 genes that were differentially regulated between the two generations (465 upregulated in the sporophyte and 702 upregulated in the gametophyte; Table supplement 3). The predicted functions of these generation-biased genes was analysed using a system of manually-assigned functional categories, together with analyses based on GO terms and KEGG pathways. The set of generation-biased genes was significantly enriched in genes belonging to two of the manually-assigned categories: “Cell wall and extracellular” and “Cellular regulation and signalling” and for genes of unknown function (Figure 3C-table supplement 3). Enriched GO terms also included several signalling- and cell wall-associated terms and terms associated with membrane transport (Figure 3D-table supplement 4). The gametophyte-biased gene set was enriched for several cell signalling KEGG pathways whereas the sporophyte-biased gene set was enriched for metabolic pathways (Figure 3E-table supplement 5). We also noted that the generation-biased genes included 23 predicted transcription factors and ten members of the EsV-1-7 domain family (Table supplement 3) (Macaisne et al., 2017). The latter were significantly enriched in the sporophyte-biased gene set (*χ*2 test p=0.001).

Both the sporophyte-biased and the gametophyte-biased datasets were enriched in genes that were predicted to encode secreted proteins (Fisher’s Exact Test p=2.02e^-8^ and p=4.14e^-6^, respectively; Table supplement 3). Analysis of GO terms associated with the secreted proteins indicated a similar pattern of enrichment to that observed for the complete set of generation-biased genes (terms associated with signalling, cell wall and membrane transport; Table supplement 4). Figure 3C illustrates the relative abundances of manually-assigned functional categories represented in the generation-biased genes predicted to encode secreted proteins.

The lists of differentially expressed genes identified by the above analysis were used to select 200 genes that showed strong differential expression between the sporophyte and gametophyte generations. The pattern of expression of the 200 genes was then analysed in the *oro* and *sam* mutants and a third mutant, *immediate upright* (*imm*), which does not cause switching between life cycle generations (Macaisne et al., 2017), as a control. Figure 3F shows that mutation of either *ORO* or *SAM* leads to upregulation of gametophyte generation genes and down-regulation of sporophyte generation genes, consistent with the switch from sporophyte to gametophyte phenotypic function. Moreover, *oro* and *sam* mutants exhibited similar patterns of expression but the patterns were markedly different to that of the *imm* mutant. Taken together with the morphological and reproductive phenotypes of the *oro* and *sam* mutants, this analysis supports the conclusion that *ORO* and *SAM* are master regulators of the gametophyte-to-sporophyte transition.

### The ORO and SAM proteins interact *in vitro*

HD TFs that act as life cycle regulators or mating type determinants often form heterodimeric complexes (Banham et al., 1995; Horst et al., 2016; Hull et al., 2005; Kämper et al., 1995; Lee et al., 2008). The ORO and SAM proteins were also shown to be capable of forming a stable heterodimer using an *in vitro* pull-down approach (Figure 4). Deletion analysis indicated that the interaction between the two proteins was mediated by their homeodomains.

**Figure 4.**
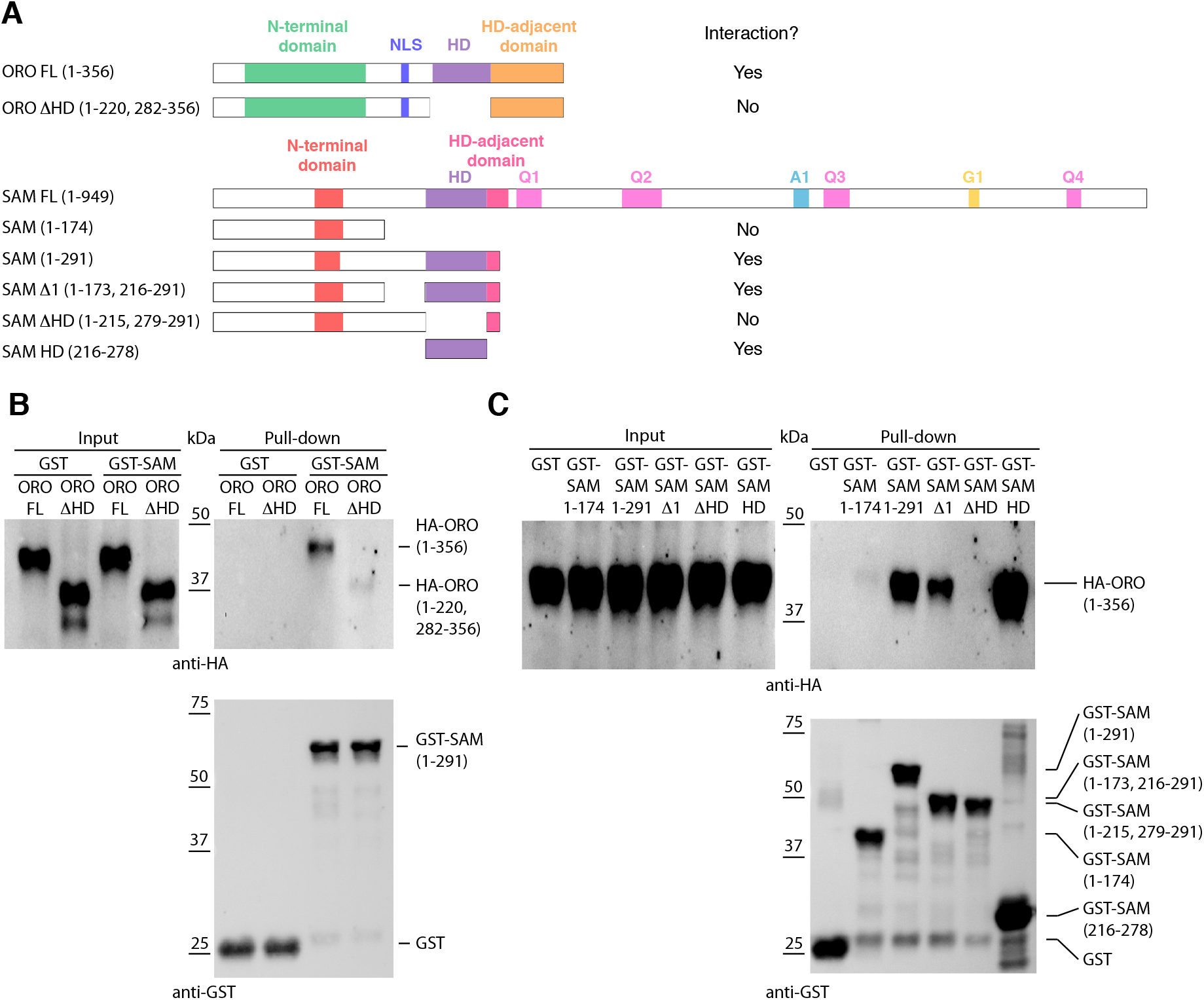
Detection of ORO-SAM heterodimerisation *in vitro* using a pull-down assay. **(A)** ORO and SAM constructs used for the pull-down experiments**. (B)** Pull-down assay between SAM and different versions of the ORO protein. **(C)** Pull-down assay between different versions of the SAM protein and full-length ORO protein. Note that all ORO proteins were fused with the HA epitope. FL, full-length; HD, homeodomain.

### Evolutionary origins and domain structure of the *ORO* and *SAM* genes

Analysis of sequence databases indicated that all brown algae possess three HD TFs, all of the TALE class, including orthologues of ORO and SAM (Figure 5A-table supplement 6). Comparison of brown algal ORO and SAM orthologues identified conserved domains both upstream and downstream of the HDs in both ORO and SAM (Figure 5B,C-figure supplement 5). These domains do not correspond to any known domains in public domain databases and were not found in any other proteins in the public sequence databases. The HD was the only domain found in both the ORO and SAM proteins (Figure 5).

**Figure 5.**
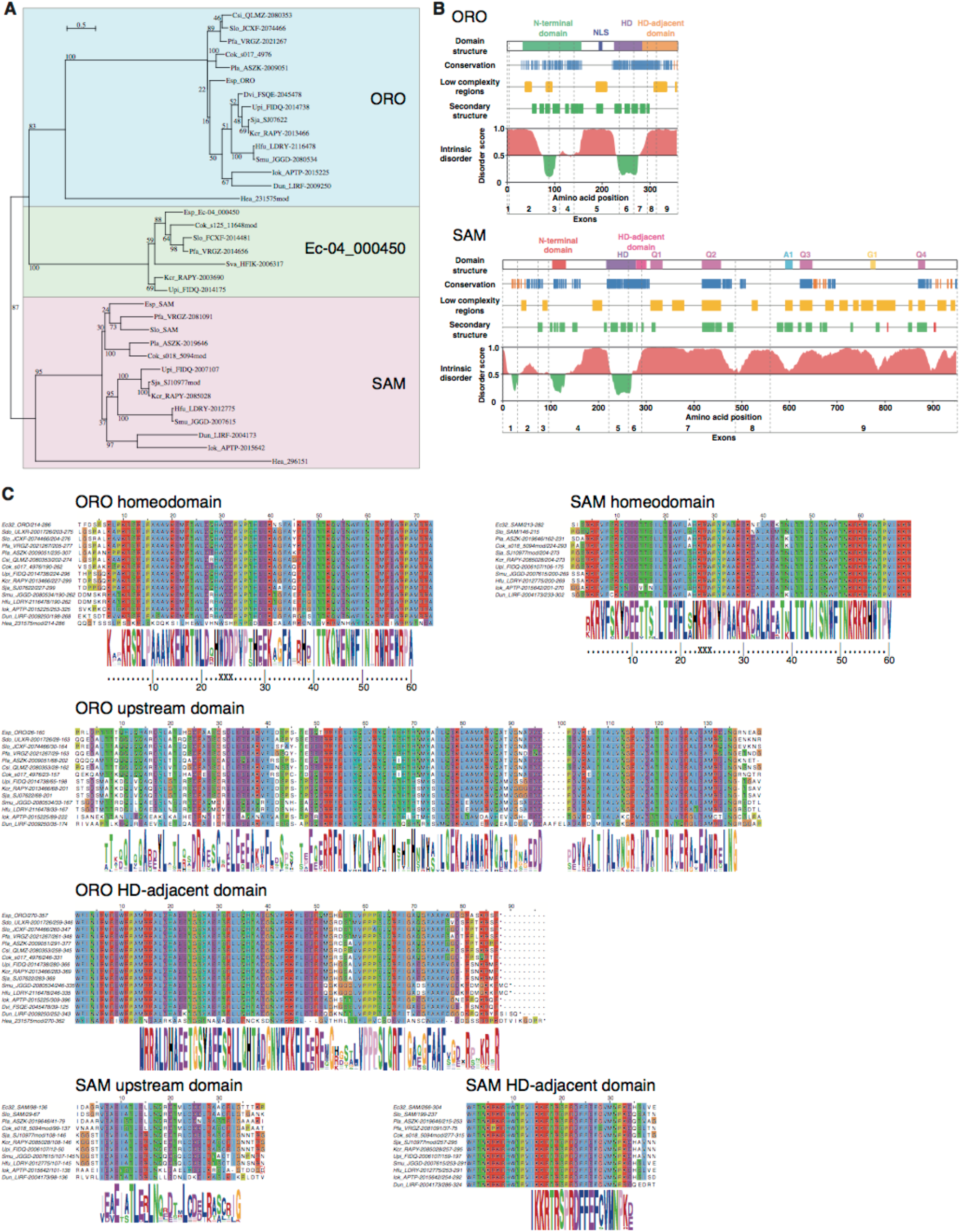
ORO and SAM conservation and domain structure. **(A)** Unrooted maximum likelihood tree of ORO, SAM and Ec-04_000450 orthologues from diverse brown algal species and the raphidophyte *Heterosigma akashiwo***. (B)** Domain structure of the ORO and SAM TALE homeodomain transcription factors. Conservation: strong (blue), less strong (orange), secondary structure: α-helix (green), β-strand (red). Q1-4, A1 and G1: regions rich in glutamine, alanine and glycine, respectively. **(C)** Conserved domains in ORO and SAM proteins. Cok, *Cladosiphon okamuranus*; Csi, *Colpomenia sinuosa*; Dvi, *Desmarestia viridis*; Dun, *Dictyopteris undulata*; Esp, *Ectocarpus* sp.; Hea, *Heterosigma akashiwo*; Hfu, *Hizikia fusiformis*; Iok, *Ishige okamurai*; Kcr, *Kjellmaniella crassifolia*; Pfa, *Petalonia fascia*; Pla, *Punctaria latifolia*; Sja, *Saccharina japonica*; Smu, *Sargassum muticum*; Sva, *Sargassum vachellianum*; Sdo, *Scytosiphon dotyi*; Slo, *Scytosiphon lomentaria*; Upi, *Undaria pinnatifida*.

To identify more distantly-related orthologues of ORO and SAM, we searched a broad range of stramenopile TALE HD TFs for the presence of characteristic ORO and SAM protein domains. Only one non-brown-algal protein, from the raphidophyte *Heterosigma akashiwo*, possessed similarity to these domains, allowing it to be classed tentatively as an ORO orthologue (gene identifier 231575mod; Figure 5A,C-table supplement 6). The transcriptome of this strain also included a truncated TALE HD TF transcript similar to SAM but more complete sequence data will be required to confirm orthology with SAM (gene identifier 296151; Figure 5A-table supplement 6). This analysis allowed the origin of ORO to be traced back to the common ancestor with the raphidophytes (about 360 Mya; Brown and Sorhannus, 2010) but the rate of divergence of the non-HD regions of ORO and SAM precluded the detection of more distantly related orthologues. An additional search based on looking for TALE HD TF genes with intron positions corresponding to those of *ORO* and *SAM* did not detect any further orthologues (Figure supplement 3).

## Discussion

The analysis presented here demonstrates that two TALE HD TFs, which are capable of forming a heterodimer, are required for the deployment of the sporophyte program during the life cycle of the brown alga *Ectocarpus*. The parallels with life cycle regulation in the green lineage, where TALE HD TFs have also been shown to regulate deployment of the sporophyte program (Horst et al., 2016; Sakakibara et al., 2013), are striking. Knockout of the KNOX class TALE HD TF genes *MKN1* and *MKN6* in *Physcomitrella patens* result in conversion of the sporophyte generation into a functional gametophyte (Sakakibara et al., 2013), essentially the same phenotype as that observed with *Ectocarpus oro* or *sam* mutants despite the fact that more than a billion years of evolution separate the two lineages (Eme et al., 2014) and that the two lineages independently evolved complex multicellularity. The similarities between life cycle regulators in the two eukaryotic supergroups suggests that they are derived from a common ancestral system that would therefore date back to early eukaryotic evolution. The ancient origin of this life cycle regulatory system is supported by the fact that distantly-related homeodomain or homeodomain-like proteins act as mating type factors in both fungi and social amoebae (Hedgethorne et al., 2017; Hull et al., 2005; Nasmyth and Shore, 1987; Van Heeckeren et al., 1998). It has been proposed that the ancestral function of this homeodomain-based life cycle regulators was to detect syngamy and to implement processes specific to the diploid phase of the life cycle such as repressing gamete formation and initiating meiosis (Perrin, 2012 and references therein). With the emergence of complex, multicellular organisms, it would not have been surprising if additional processes such as developmental networks had come under the control of these regulators as this would have ensured that those developmental processes were deployed at the appropriate stage of the life cycle (Cock et al., 2013). Indeed, it has been suggested that modifications to homeodomain-based regulatory circuits may have played an important role in the emergence of sporophyte complexity in the green lineage (Bowman et al., 2016; Lee et al., 2008). Key events may have included the replacement of the Gsp1-like class of BELL-related1 genes with alternative (true BEL-class) proteins and diversification of both the true BELL-class and the KNOX-class TALE HD TFs. In particular, the emergence and subfunctionalisation of two KNOX subfamilies early in streptophyte evolution is thought to have facilitated the evolution of more complex sporophyte transcriptional networks (Furumizu et al., 2015; Sakakibara et al., 2013). In the brown algae, ORO and SAM also function as major developmental regulators but, in this lineage, the emergence of a multicellular sporophyte has not been associated with a marked expansion of the TALE HD TF family. However, there does appear to have been considerable divergence of the ORO and SAM protein sequences during brown algal evolution, perhaps reflecting the evolution of new functions associated with multicellular development and divergence of the sporophyte and gametophyte developmental programs. Heterodimerisation appears to be a conserved feature of brown algal and green lineage TALE HD TFs (Figure 4 and Lee et al., 2008) despite the lack of domain conservation. However, in *Ectocarpus* heterodimerisation involves the ORO and SAM HDs whereas in *Chlamydomonas*, it is the KNOX1 and KNOX2 domains of Gsm1 that interact with the C-terminal region of Gsp1 (which includes the HD, Ala and DE domains).

Interestingly, diploid sporophytes heterozygous for *sam* mutations exhibited abortive development of unilocular sporangia at a stage corresponding to the meiotic division of the mother cell. At first sight it might seem surprising that a gene should play an important role both directly following the haploid to diploid transition (initiation of sporophyte development) and at the opposite end of the life cycle, during the diploid to haploid transition (meiosis). However, these phenotypes make more sense when viewed from an evolutionary perspective, if the ORO SAM system originally evolved as a global regulator of diploid phase processes.

There is now accumulating evidence for an ancient role for HD TFs in life cycle regulation in both the bikont and unikont branches of the eukaryotic tree of life (Hedgethorne et al., 2017; Horst et al., 2016; Hull et al., 2005; Lee et al., 2008; Sakakibara et al., 2013 and this study). We show here that these systems have been adapted to coordinate life cycle progression and development in at least two multicellular eukaryotic lineages (land plants and brown algae). The recruitment of TALE HD TFs as sporophyte program master regulators in both the brown and green lineages represents a particularly interesting example of latent homology, where the shared ancestral genetic toolkit constrains the evolutionary process in two diverging lineages leading to convergent evolution of similar regulatory systems (Nagy et al., 2014). The identification of such constraints through comparative analysis of independent complex multicellular lineages provides important insights into the evolutionary processes underlying the emergence of complex multicellularity. One particularly interesting outstanding question is whether HD TFs also play a role in coordinating life cycle progression and development in animals? Analysis of the functions of TALE HD TFs in unicellular relatives of animals may help provide some insights into this question.

## Materials and Methods

### Treatment with the sporophyte-produced diffusible factor

Sporophyte-conditioned medium, gametophyte-conditioned medium and protoplasts were produced as previously described (Arun et al., 2013). Protoplasts were allowed to regenerate either in sporophyte-conditioned medium supplemented with osmoticum or in gametophyte-conditioned supplemented with osmoticum as a control. Congo red staining was used to distinguish sporophytes from gametophytes (Arun et al., 2013). At least 60 individuals were scored per treatment per experiment. Results are representative of three independent experiments.

### Mapping of genetic loci

The *oro* mutation has been shown to behave as a single-locus, recessive, Mendelian factor (Coelho et al., 2011). AFLP analysis was carried out essentially as described by Vos et al. (1995). DNA was extracted from 50 wild type and 50 *oro* individuals derived from a cross between the outcrossing line Ec568 (Heesch et al., 2010) and the *oro* mutant Ec494 (Coelho et al., 2011; Table supplement 1). Equal amounts of DNA were combined into two pools, for bulk segregant analysis. Pre-selective amplification was carried out with an *Eco*RI-anchored primer and an *Mse*I-anchored primer, each with one selective nucleotide, in five different combinations (*Eco*RI+T / *Mse*I+G; *Eco*RI+T / *Mse*I+A; *Eco*RI+C / *Mse*I+G; *Eco*RI+C / *Mse*I+A; *Eco*RI+A / *Mse*I+C). These reactions were diluted 1:150 for the selective amplifications. The selective amplifications used an *Eco*RI-anchored primer and an *Mse*I-anchored primer, each with three selective nucleotides, in various different combinations. The PCR conditions for both steps were 94°C for 30 sec, followed by 20 cycles of DNA amplification (30 sec at 94°C, 1 min at 56°C and 1 min at 72°C) and a 5 min incubation at 72°C except that this protocol was preceded by 13 touchdown cycles involving a decrease of 0.7°C per cycle for the selective amplifications. PCR products were analysed on a LI-COR apparatus. This analysis identified two flanking AFLP markers located at 20.3 cM and 21.1 cM on either side of the *ORO* locus. For 23 (12 *oro* and 11 wild type) of the 100 individuals, no recombination events were detected within the 41.4 cM interval between the two markers. Screening of these 23 individuals (11 wild type and 12 *oro*) with the microsatellite markers previously developed for a sequence-anchored genetic map (Heesch et al., 2010) identified one marker within the 41.4 cM interval (M_512) and located the *ORO* locus to near the bottom of chromosome 14 (Cormier et al., 2017).

Fine mapping employed a segregating population of 2,000 individuals derived from the cross between the *oro* mutant line (Ec494) and the outcrossing line Ec568 and an additional 11 microsatellite markers within the mapping interval (Table supplement 7) designed based on the *Ectocarpus* genome sequence (Cock et al., 2010). PCR reactions contained 5 ng of template DNA, 1.5 μl of 5xGoTaq reaction buffer, 0.25 units of GoTaq-polymerase (Promega), 10 nmol MgCl_2_, 0.25 μl of dimethyl sulphoxide, 0.5 nmol of each dNTP, 2 pmol of the reverse primer, 0.2 pmol of the forward primer (which included a 19-base tail that corresponded to a nucleotide sequence of the M13 bacteriophage) and 1.8 pmol of the fluorescence marked M13 primer. The PCR conditions were 94°C for 4 min followed by 13 touch-down cycles (94°C for 30 sec, 65-54°C for 1 min and 72°C for 30 sec) and 25 cycles at 94°C for 30 sec, 53°C for 1 min and 72°C for 30 sec. Samples were genotyped by electrophoresis on an ABI3130xl Genetic Analyser (Applied Biosystems) and analysis with Genemapper version 4.0 (Applied Biosystems). Using the microsatellite markers, the *oro* mutation was mapped to a 34.5 kbp (0.45 cM) interval, which contained five genes. Analysis of an assembled, complete genome sequence for a strain carrying the *oro* mutation (strain Ec597; European Nucleotide Archive PRJEB1869; Ahmed et al., 2014) together with Sanger method resequencing of ambiguous regions demonstrated that there was only one mutation within the mapped interval: an 11 bp deletion in the gene with the LocusID Ec-14_005920.

### Reconstruction and sequence correction of the *ORO* and *SAM* loci

The sequence of the 34.5 kbp mapped interval containing the *ORO* gene (chromosome 27, 5463270-5497776) in the wild type *Ectocarpus* reference strain Ec32 included one short region of uncertain sequence 1026 bp downstream of the end of the *ORO* open reading frame. The sequence of this region was completed by PCR amplification and Sanger sequencing and confirmed by mapping Illumina read data to the corrected region. The corrected *ORO* gene region has been submitted to Genbank under the accession number KU746822.

Comparison of the reference genome (strain Ec32) supercontig that contains the *SAM* gene (sctg_251) with homologous supercontigs from several independently assembled draft genome sequences corresponding to closely related *Ectocarpus* sp. strains (Ahmed et al., 2014; Cormier et al., 2017) indicated that sctg_251 was chimeric and that the first three exons of the *SAM* gene were missing. The complete *SAM* gene was therefore assembled and has been submitted to Genbank under the accession number KU746823.

### Quantitative reverse transcriptase polymerase chain reaction analysis of mRNA abundance

Total RNA was extracted from wild-type gametophytes and partheno-sporophytes (Ec32) and from *sam-1* (Ec374) and *sam-2* (Ec364) partheno-gametophytes using the Qiagen RNeasy Plant mini kit and any contaminating DNA was removed by digestion with Ambion Turbo DNase (Life Technologies). The generation marker genes analysed were Ec-20_001150 and Ec-26_000310 (sporophyte markers), and Ec-23_004240 and Ec-21_006530 (gametophyte markers), which are referred to as *IDW6, IDW7, IUP2* and *IUP7* respectively, in Peters *et al*. (2008). Following reverse transcription of 50-350 ng total RNA with the ImPro II TM Reverse Transcription System (Promega), quantitative RT-PCR was performed on LightCycler^®^ 480 II instrument (Roche). Reactions were run in 10 μl containing 5 ng cDNA, 500nM of each oligo and 1x LightCycler^®^ 480 DNA SYBR Green I mix (Roche). The sequences of the oligonucleotides used are listed in Table supplement 8. Pre-amplification was performed at 95°C for 5 min, followed by the amplification reaction consisting of 45 cycles of 95°C for 10 sec, 60°C for 30 sec and 72°C for 15 sec with recording of the fluorescent signal after each cycle. Amplification specificity and efficiency were checked using a melting curve and a genomic DNA dilution series, respectively, and efficiency was always between 90% and 110%. Data were analysed using the LightCycler^®^ 480 software (release 1.5.0). A pair of primers that amplified a fragment which spanned intron 2 of the *SAM* gene was used to verify that there was no contaminating DNA (Table supplement 8). Standard curves generated from serial dilutions of genomic DNA allowed quantification for each gene. Gene expression was normalized against the reference gene *EEF1A2*. Three technical replicates were performed for the standard curves and for each sample. Statistical analysis (Kruskal-Wallis test and Dunn’s Multiple Comparison Post Test) was performed using the software GraphPadPrism5.

### RNA-seq analysis

RNA for RNA-seq analysis was extracted from duplicate samples (two biological replicates) of approximately 300 mg (wet weight) of tissue either using the Qiagen RNeasy plant mini kit with an on-column Deoxyribonuclease I treatment or following a modified version (Peters et al., 2008) of the protocol described by Apt et al. (1995). Briefly, this second protocol involved extraction with a cetyltrimethylammonium bromide (CTAB)-based buffer and subsequent phenol-chloroform purification, LiCl-precipitation, and DNAse digestion (Turbo DNAse, Ambion, Austin, TX, USA) steps. RNA quality and concentration was then analysed on 1.5% agarose gel stained with ethidium bromide and a NanoDrop ND-1000 spectrophotometer (NanoDrop products, Wilmington, DE, USA). Between 21 and 93 million sequence reads were generated for each sample on an Illumina Hi-seq2000 platform (Table supplement 9). Raw reads were quality trimmed with Trimmomatic (leading and trailing bases with quality below 3 and the first 12 bases were removed, minimum read length 50 bp) (Bolger et al., 2014). High score reads were aligned to the *Ectocarpus* reference genome (Cock et al., 2010; available at Orcae; Sterck et al., 2012) using Tophat2 with the Bowtie2 aligner (Kim et al., 2013). The mapped sequencing data was then processed with HTSeq (Anders et al., 2014) to obtain counts for sequencing reads mapped to exons. Expression values were represented as TPM and TPM>1 was applied as a filter to remove noise.

Differential expression was detected using the DESeq2 package (Bioconductor; Love et al., 2014) using an adjusted p-value cut-off of 0.05 and a minimal fold-change of two. Heatmaps were generated using the Heatplus package for R (Ploner, 2015) and colour schemes selected from the ColorBrewer project (http://colorbrewer.org).

The entire set of 16,724 protein-coding genes in the *Ectocarpus* Ec32 genome were manually assigned to one of 22 functional categories (Table supplement 10) and this information was used to determine whether sets of differentially expressed genes were enriched in particular functional categories compared to the entire nuclear genome (*χ*^2^ test). Blast2GO (Conesa and Götz, 2008) was used to detect enrichment of GO-terms associated with the genes that were consistently up- or downregulated in pairwise comparisons of the wild type gametophyte, the *sam* mutant and the *oro* mutant with the wild type sporophyte. Significance was determined using a Fisher exact test with an FDR corrected p-value cutoff of 0.05. Sub-cellular localisations of proteins were predicted using Hectar (Gschloessl et al., 2008). Sets of secreted proteins corresponded to those predicted to possess a signal peptide or a signal anchor.

### Detection of protein-protein interactions

Pull-down assays were carried out using the MagneGST™ Pull-Down System (Promega, Madison, WI) by combining human influenza hemagglutinin (HA)-tagged and glutathione S-transferase (GST) fusion proteins. *In vitro* transcription/translation of HA-tagged ORO proteins was carried out using the TNT^®^ Coupled Wheat Germ Extract System (Promega, Madison, WI). GST-tagged SAM proteins were expressed in *Escherichia coli.* Protein production was induced by adding IPTG to a final concentration of 2mM and shaking for 20 h at 16°C. After the capture phase, beads were washed four times with 400 μL of washing buffer (0.5% IGEPAL, 290 mM NaCl, 10 mM KCl, 4.2 mM Na_2_HPO_4_, 2 mM KH_2_PO_4_, at pH 7.2) at room temperature. Beads were then recovered in SDS-PAGE loading buffer, and proteins analysed by SDS-PAGE followed by Clarity™ chemiluminescent detection (Biorad, Hercules, CA). The anti-HA antibody (3F10) was purchased from Roche, and the anti-GST antibody (91G1) from Ozyme.

### Searches for HD proteins from other stramenopile species

Searches for homeodomain proteins from additional brown algal or stramenopile species were carried out against the NCBI, Uniprot, oneKP (Matasci et al., 2014) and iMicrobe databases and against sequence databases for individual brown algal (*Saccharina japonica,* Ye et al., 2015; *Cladosiphon okamuranus,* Nishitsuji et al., 2016) and stramenopile genomes

(*Nannochloropsis oceanica, Aureococcus anophagefferens, Phaeodactylum tricornutum, Thalassiosira pseudonana, Pseudo-nitzschia multiseries*) and transcriptomes (*Vaucheria litorea, Heterosigma akashiwo*) using both Blast (Blastp or tBlastn) and HMMsearch with a number of different alignments of brown algal TALE HD TF proteins. As the homeodomain alone does not provide enough information to construct well-supported phylogenetic trees, searches for ORO and SAM orthologues were based on screening for the presence of the additional protein domains conserved in brown algal ORO and SAM proteins.

As intron position and phase was strongly conserved between the homeoboxes of *ORO* and *SAM* orthologues within the brown algae, this information was also used to search for ORO and SAM orthologues in other stramenopile lineages. However, this analysis failed to detect any additional candidate *ORO* or *SAM* orthologues. These observations are consistent with a similar analysis of plant homeobox introns, which showed that intron positions were strongly conserved in recently diverged classes of homeobox gene but concluded that homeobox introns were of limited utility to deduce ancient evolutionary relationships (Mukherjee et al., 2009).

GenomeView (Abeel et al., 2012) was used together with publically available genome and RNA-seq sequence data (Nishitsuji et al., 2016; Ye et al., 2015) to improve the gene models for some of the brown algal TALE HD TFs (indicated in Table supplement 6 by adding the suffix “mod” for modified to the protein identifier).

### Phylogenetic analysis and protein analysis and comparisons

Multiple alignments were generated with Muscle in MEGA7 (Tamura et al., 2011). Phylogenetic trees were then generated with RAxML (Stamatakis, 2015) using 1000 bootstrap replicates and the most appropriate model based on an analysis in MEGA7. Domain alignments were constructed in Jalview (http://www.jalview.org/) and consensus sequence logos were generated with WebLogo (http://weblogo.berkeley.edu/logo.cgi). Intrinsic disorder in protein folding was predicted using SPINE-D (Zhang et al., 2012), low complexity regions with SEG (default parameters, 12 amino acid window; Wootton, 1994) and secondary structure with PSIPRED (Buchan et al., 2013).

## Acknowledgements

We thank the ABiMS platform (Roscoff Marine Station) for providing computing facilities and support.

## Additional information

### Author contributions

S.M.C., O.G., D.S. and A.F.P. isolated life cycle mutants and carried out culture work. A.A., S.M.C., A.F.P., D.S., C.T. and A.B. performed the positional cloning. L.P. and S.B. analysed protein interactions. H.Y. and S.M.C. carried out diffusible factor experiments. M.S., G.J.M., N.M. and D.S. generated expression and sequence data. A.P.L., K.A., S.M.C. and J.M.C. analysed data. J.M.C. designed and supervised the research and wrote the article with help from all the authors.

## Additional files

### Supplementary files

#### Supplementary notes

Supplementary Figure 1. Morphological characteristics and response to unidirectional light of *sam* mutants.

Supplementary Figure 2. Evidence for the production of full-length *ORO* and *SAM* transcripts during the gametophyte generation.

Supplementary Figure 3. Intron conservation in homeobox genes.

Supplementary Table 1. *Ectocarpus* strains used in this study.

Supplementary Table 2. Congo red staining of wild type or *sam-1* protoplasts following regeneration in sporophyte-conditioned medium (SCM) or gametophyte-conditioned medium (GCM).

Supplementary Table 3. Analysis of genes that are differentially expressed in the gametophyte and sporophyte generations.

Supplementary Table 4. Gene ontology analysis of the gametophyte versus sporophyte differentially regulated genes.

Supplementary Table 5. Kyoto encyclopaedia of genes and genomes (KEGG) pathway analysis of the gametophyte versus sporophyte differentially regulated genes.

Supplementary Table 6. TALE homeodomain transcription factors in brown algae and other stramenopiles.

Supplementary Table 7. New microsatellite markers developed to map the *ORO* gene.

Supplementary Table 8. Oligonucleotides used for the qRT-PCR analysis.

Supplementary Table 9. *Ectocarpus* RNA-seq data used in this study.

Supplementary Table 10. Manual functional assignments and Hectar subcellular targeting predictions for all *Ectocarpus* nucleus-encoded proteins

## Supplementary information

### Supplementary notes

#### Expression of *ORO* and *SAM* during the gametophyte generation

Gametophytes carrying *oro* or *sam* mutations did not exhibit any obvious phenotypic defects, despite the fact that both genes are expressed during this generation (although *SAM* expression was very weak). In *P. patens,* GUS fusion experiments failed to detect expression of KNOX genes in the gametophyte but RT-PCR analysis and cDNA cloning has indicated that KNOX (and BEL) transcripts are expressed during this generation (Champagne and Ashton, 2001; Sakakibara et al., 2013, 2008). However, no phenotypes were detected during the haploid protonema or gametophore stages in KNOX mutant lines (Sakakibara et al., 2013, 2008;, Singer and Ashton, 2007) and the RT-PCR only amplified certain regions of the transcripts. Consequently, these results have been interpreted as evidence for the presence of partial transcripts during the gametophyte generation. To determine whether the *ORO* and *SAM* transcripts produced in *Ectocarpus* were incomplete, RNA-seq data from male and female, immature and mature gametophytes was mapped onto the *ORO* and *SAM* gene sequences. This analysis indicated that full-length transcripts of both the *ORO* and *SAM* genes are produced during the gametophyte generation (Figure supplement 2).

#### ORO and SAM domain structure

The conserved domains that flank the homeodomains in the ORO and SAM proteins share no detectable similarity with domains that are associated with TALE HDs in the green (Viridiplantae) lineage, such as the KNOX, ELK and BEL domains. Interestingly, both the ORO and SAM proteins possess regions that are predicted to be highly disordered (Figure supplement 5B). Intrinsically disordered region are a common feature in transcription factors and the flexibility conferred by these regions is thought to allow them to interact with a broad range of partners (Niklas et al., 2015), a factor that may be important for master developmental regulators such as the ORO and SAM proteins.

#### Stramenopile TALE HD TFs

All the stramenopile species analysed in this study possessed at least two TALE HD TFs, with some species possessing as many as 14 (Table supplement 6). Note that genomes of several diverse stramenopile lineages outside the brown algae were predicted to encode proteins with more than one HD (Table supplement 6). It is possible that these proteins have the capacity to bind regulatory sequences in a similar manner to heterodimers of proteins with single HDs

#### Supplementary figures

**Figure S1.**
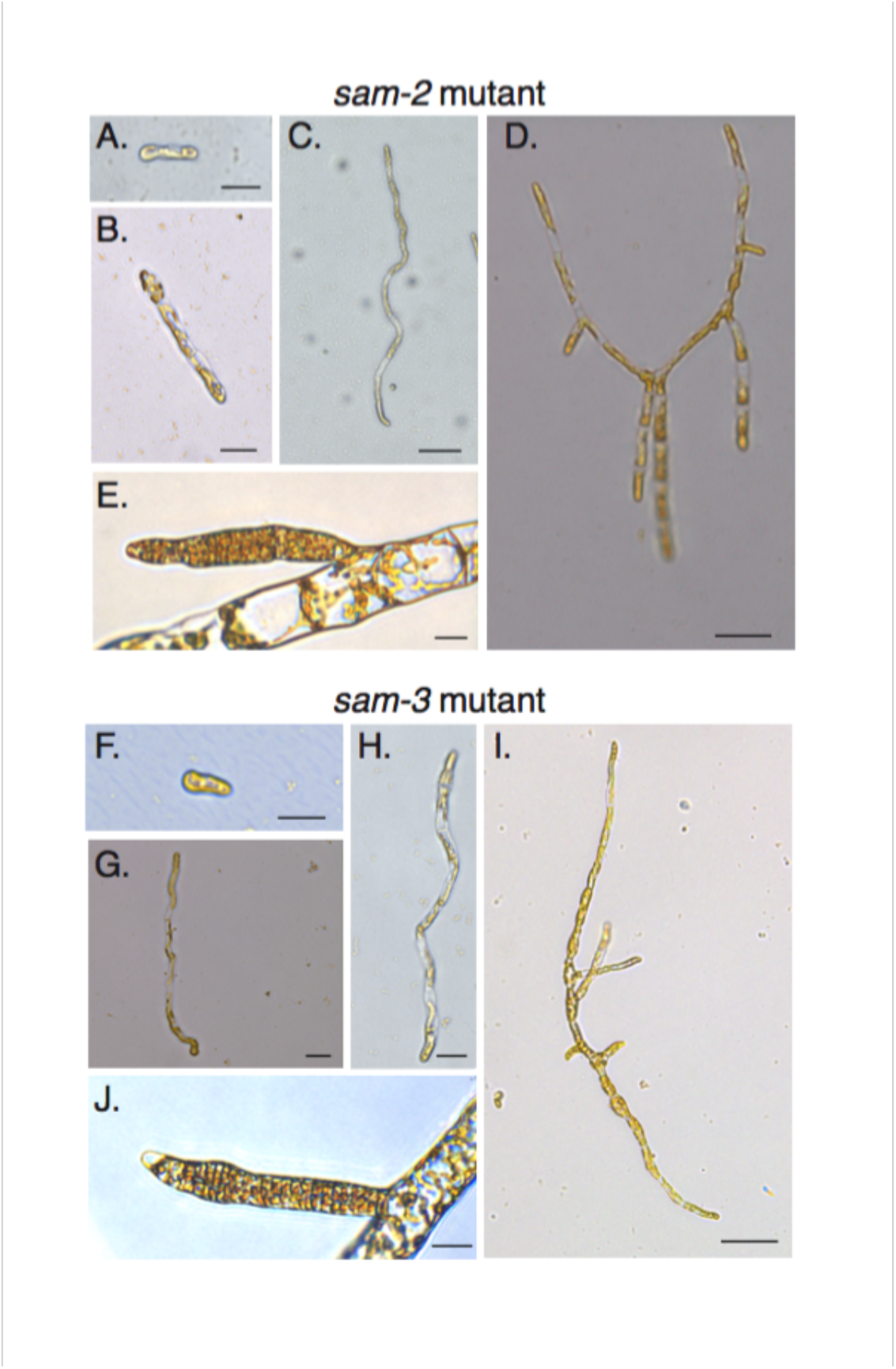
Morphological characteristics and response to unidirectional light of *sam* mutants. **(A-J)** The *sam-2* and *sam-3* mutants exhibit gametophyte-like morphological characteristics. A-E, *sam-2* mutant (strain Ec364); F-J, *sam-3* mutant (strain Ec793); A-D and F-I, different stages of early development from germination to young, branched germling; E and J, plurilocular gametangia. Size bars indicate 20 μm for all panels except C, D and I where the size bar indicate 50 μm. See Figure 2 for the equivalent developmental stages of wild type sporophytes and gametophytes.

**Figure S2.**
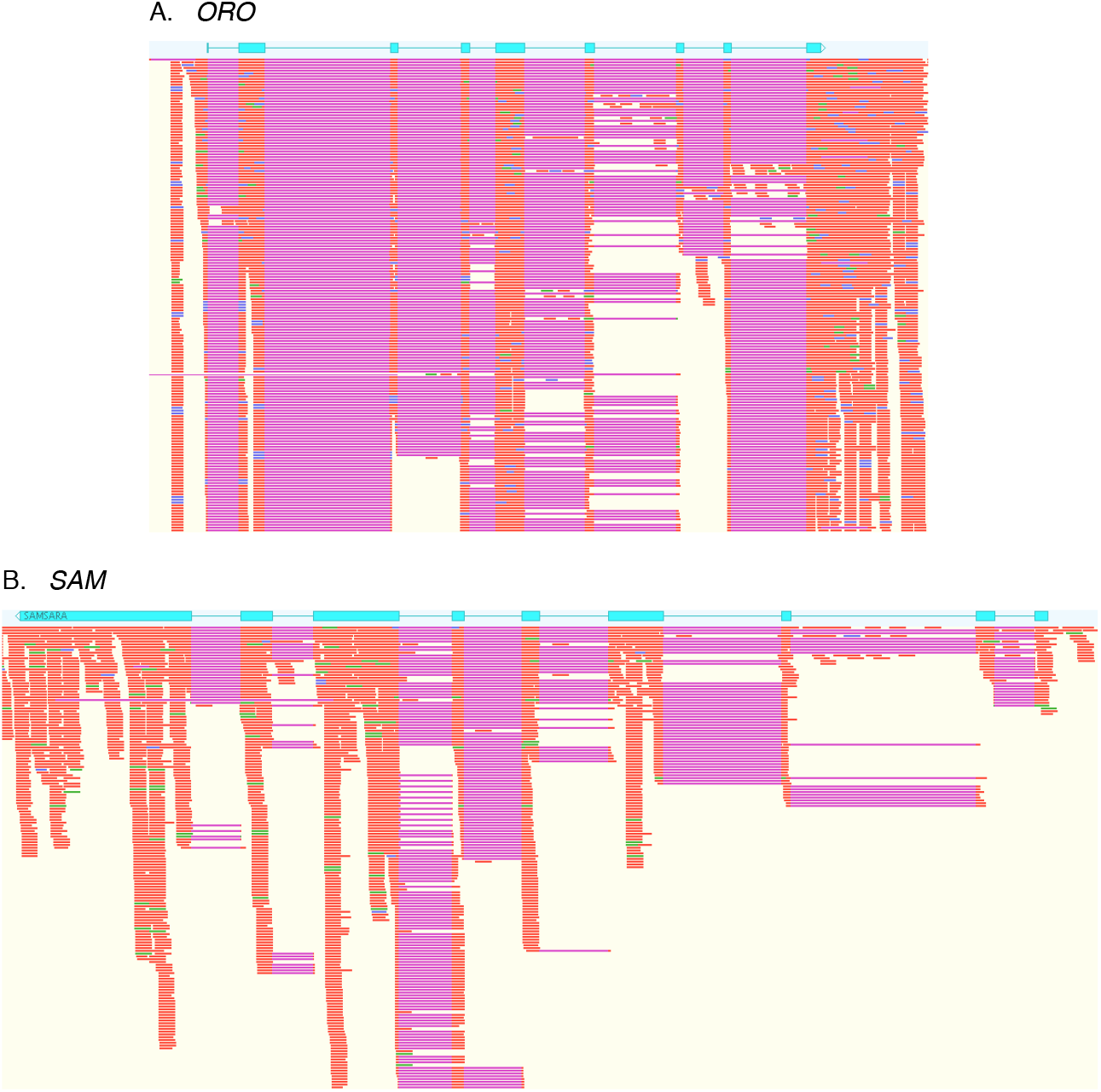
Evidence for the production of full-length *ORO* and *SAM* transcripts during the gametophyte generation. Immature and mature male and female gametophyte Illumina RNA-seq data was mapped onto the *ORO* and *SAM* gene sequences using Tophat2. Blue boxes, *ORO* and *SAM* coding exons; orange, RNA-seq reads; purple, gaps introduce during mapping corresponding to introns.

**Figure S3.**
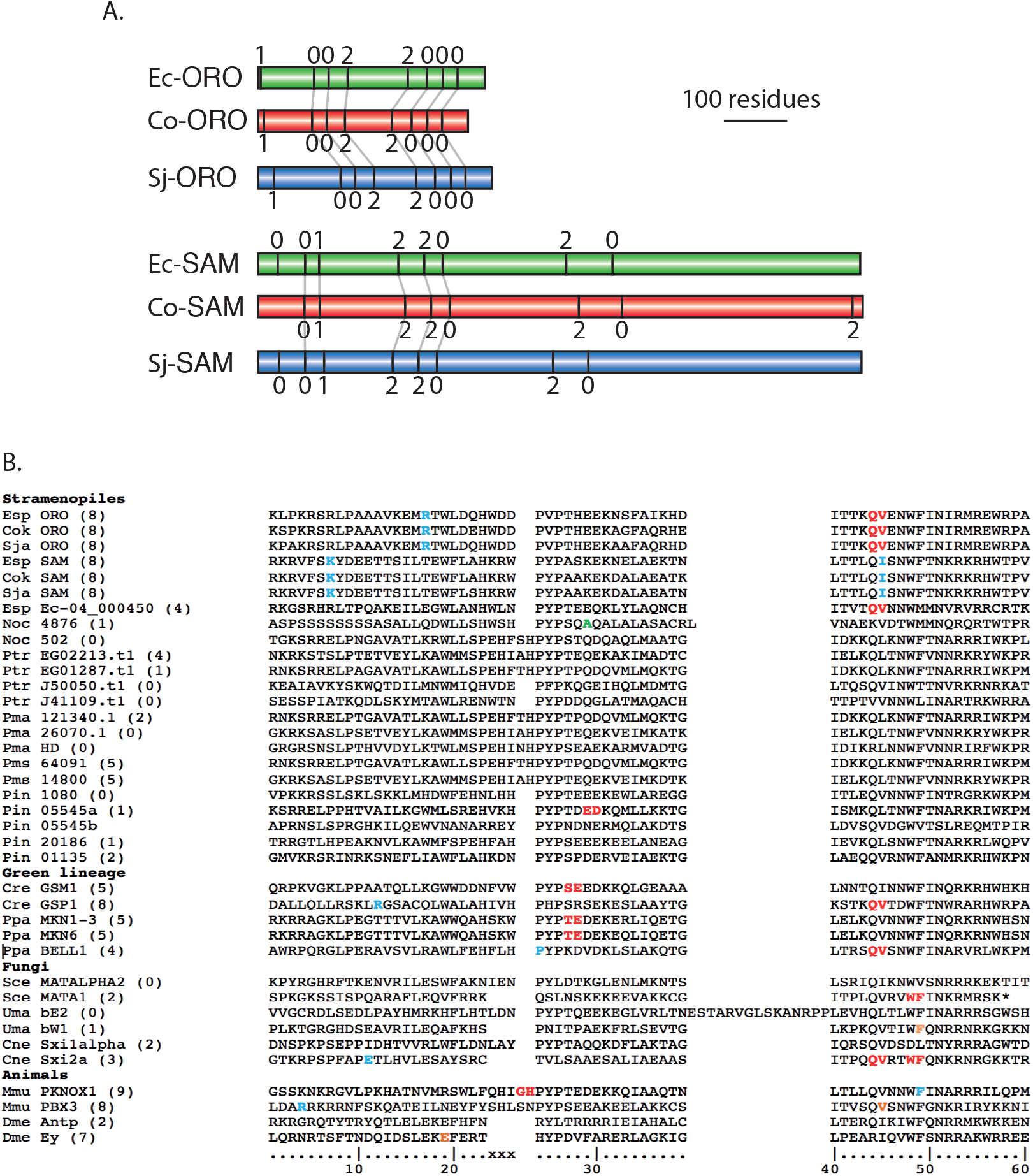
Intron conservation in homeobox genes. **(A)** Conservation of introns in *Ectocarpus* (Ec), *C. okamuranus* (Co) and *S. japonica* (Sj) *ORO* and *SAM* genes. Schematic representation of the coding regions of *ORO* and *SAM* genes showing the positions and phase of introns. Conserved intron positions, based on sequence similarity, are indicated by grey lines. Intron boundaries at similar positions but not linked by a grey line are also likely to be ancestral but it is not possible to verify homology because these regions of the proteins are too diverged. Protein identifiers are Ec-*ORO*, Ec-14_005920; Co-*ORO*, Cok_S_s017_4976.t2; Sj-*ORO*, SJ07622; Ec-*SAM*, Ec-27_006660; Co-*SAM*, Cok_S_s018_5094mod; Sj-*SAM*, SJ10977mod where the suffix "mod" indicates that the original gene model has been modified (see Table supplement 6). **(B)** Positions of homeobox introns in stramenopile homeobox genes, life cycle regulators from the green lineage, fungal mating type regulators and selected metazoan homeobox genes. Intron positions are colour coded according to phase: 0, red; 1, blue; 2, orange. The numbering at the bottom indicate the conserved 60 residues of the homeodomain and xxx indicates the three additional amino acids in TALE HD TFs. Numbers in brackets indicate total number of introns in the coding region. The asterisk indicates a stop codon. Esp, *Ectocarpus* sp.; Cok, *Cladosiphon okamuranus*; Sja, *Saccharina japonica*; Noc, *Nannochloropsis oceanica*; Ptr, *Phaeodactylum tricornutum*; Pmu, *Pseudo-nitzschia multiseries*; Cre, *Chlamydomonas reinhardtii*; Ppa, *Physcomitrella patens*; Sce, *Saccharomyces cerevisiae*; Uma, *Ustilago maydis*; Cne, *Cryptococcus neoformans*; Dme, *Drosophila melanogaster*. Note that *Phytophthora infestans* gene 05545 has two homeoboxes.

#### Supplementary tables

See separate Excel file.

